# A distributed neural architecture of sustained affect across external and internal experience

**DOI:** 10.64898/2026.05.05.723090

**Authors:** Ran Zhang, Debo Dong, Xianyang Gan, Sarah Genon, Simon B. Eickhoff, Ting Xu, Bing Cao, Qinghua He, Tingyong Feng, Benjamin Becker, Feng Zhou

## Abstract

Sustained affect shapes well-being, yet its neural architecture across externally elicited and internally generated experience remains unclear. Using whole-brain functional connectivity during minutes-long naturalistic movie viewing, we derived positive and negative affective experience signatures and their underlying neural architecture. These signatures predicted valence-specific affective intensity and generalized to independent movie-viewing data and internally generated affect, discriminating sad memory and rumination from neutral distraction while tracking subjective experience. Importantly, their expression showed little relation to vigilance or cognitive demand. Characterization of these signatures revealed coherent community structure and a shared distributed backbone, alongside valence-preferential components, consistent with a partially separable architecture. Extending beyond experimentally evoked states, in four resting-state depression cohorts, these signatures distinguished patients from controls with reduced positive and elevated negative signature expression, and predicted symptom burden and anhedonia. These findings identify a generalizable distributed architecture bridging external and internal affective experience and extending to clinically relevant affective dysregulation.

## Introduction

Affective experience—whether triggered by external events or internally generated thoughts—is a fundamental determinant of behavior and well-being^1,2^. When these affective states become persistently maladaptive, they manifest as hallmark features of mental disorders, such as sustained negative mood and anhedonia in depression^3,4^. Yet objective quantification of sustained affective experience remains challenging. Subjective feelings can dissociate from physiological and behavioral responses^5–7^, leaving their assessment in laboratory and clinical settings heavily reliant on self-report, which can be unreliable^8^ or infeasible in vulnerable populations. Neurobiological markers derived from brain features may complement self-report and help clarify how distributed brain systems support affective experience^9^.

To date, most multivariate neuromarkers for subjective affective experience target acute, stimulus-locked responses to simplified cues (e.g., pictures) lasting only a few seconds^5,6,10–14^. However, the neural dynamics supporting phasic affect may differ from those underlying sustained affect^15–17^, which unfolds over minutes and longer timescales, for example during movie viewing or during internally generated affective states (e.g., rumination). Consequently, transient reactivity markers are often insufficient to capture these extended states^17,18^. This gap is particularly consequential for clinical translation, because affective disorders are often characterized by persistent mood dysregulation.

Affective experience can be driven by external events or internally generated by memory and self-referential thought, posing an additional challenge for understanding the neural basis of sustained affect. These modes recruit partly dissociable systems^19^, raising a key question: can neural representations derived from externally elicited affect generalize to internally generated experience? This question is important not only for understanding whether sustained affect reflects a common large-scale architecture across percept-driven and self-generated states, but also for translational applications, where maladaptive affect can be stimulus-independent and self-generated^20–22^. Moreover, direct resting-state case-control decoding may show limited cross-cohort transportability, particularly at small to moderate sample sizes, motivating approaches that derive signatures from experimentally constrained affective processes rather than from diagnosis alone^23^.

Naturalistic movie viewing offers a useful platform to address these challenges, as it continuously engages dynamic multisensory integration, narrative comprehension, and higher-order appraisal over extended time windows^24–27^. This sustained engagement may better approximate the temporal dynamics of affect during self-generated thought, which unfolds within an ongoing stream of cognition rather than as discrete, stimulus-locked events^28^. Because these experiences unfold over extended timescales and rely on distributed interactions across systems, whole-brain functional connectivity (FC) provides a natural representation for modeling sustained affect^17,18,29,30^. Movie-evoked sustained affect may also engage integrative and appraisal-related substrates implicated in internally generated affect^31^, making movie viewing a useful context for identifying generalizable signatures of sustained experience.

Beyond generalization across elicitation modes, the large-scale neural organization of sustained positive and negative affect remains unclear. Dimensional theories often treat valence as a single bipolar continuum, such that increasing positivity necessarily implies decreasing negativity, with neutral experience anchored at the midpoint^32,33^. Within this framework, neural representations of positive and negative affect should largely be expressed as sign-inverted variants of a common pattern^34^. However, other accounts propose a partially separable organization in which positive and negative affect rely on partially distinct appraisals and regulatory mechanisms^35,36^, even if subjective experience is often described along a single positive-to-negative dimension. Distinguishing these possibilities requires neuromarkers that track the intensity of one valence without being driven by the opposite valence. Such valence specificity may also help mitigate confounds from valence-invariant state factors, such as vigilance or cognitive demand, which can covary with affective task conditions and influence large-scale networks, including the default mode (DMN) and frontoparietal (FPN) networks, central to affective experience^37–40^. Such confounding influences are a known limitation of many existing affective neuromarkers^7^.

Here, we developed whole-brain FC-based models to predict minutes-long positive and negative affective intensity during naturalistic movie viewing. Following previous studies^41,42^, we used affective arousal ratings—defined as how strongly an affective experience is felt, conceptually separable from other arousal-related constructs such as vigilance (wakefulness)^39^—to index the magnitude of positive and negative affect. By prioritizing within-valence intensity variation without forcing strong separation between neutral and opposite-valence conditions, we aimed to derive valence-specific representations. This enabled us to empirically test whether positive and negative affective experiences are supported by shared or partially separable macroscale networks. Across nine independent datasets, we asked whether signatures derived from externally evoked sustained affect (i) generalize to internally generated affective experience, (ii) are not attributable to valence-nonspecific confounds, including vigilance (sleep-wake state) and cognitive demand (working-memory load), (iii) reveal a shared distributed backbone across positive and negative affective experience, or whether they additionally feature valence-preferential components selectively informative for positive versus negative affect, and (iv) extend to task-free affective dysregulation in depression by differentiating patients from healthy controls in opposing directions (i.e., reduced positive and heightened negative signature expression) and tracking symptom burden and anhedonia (Fig. 1).

**Fig. 1.**
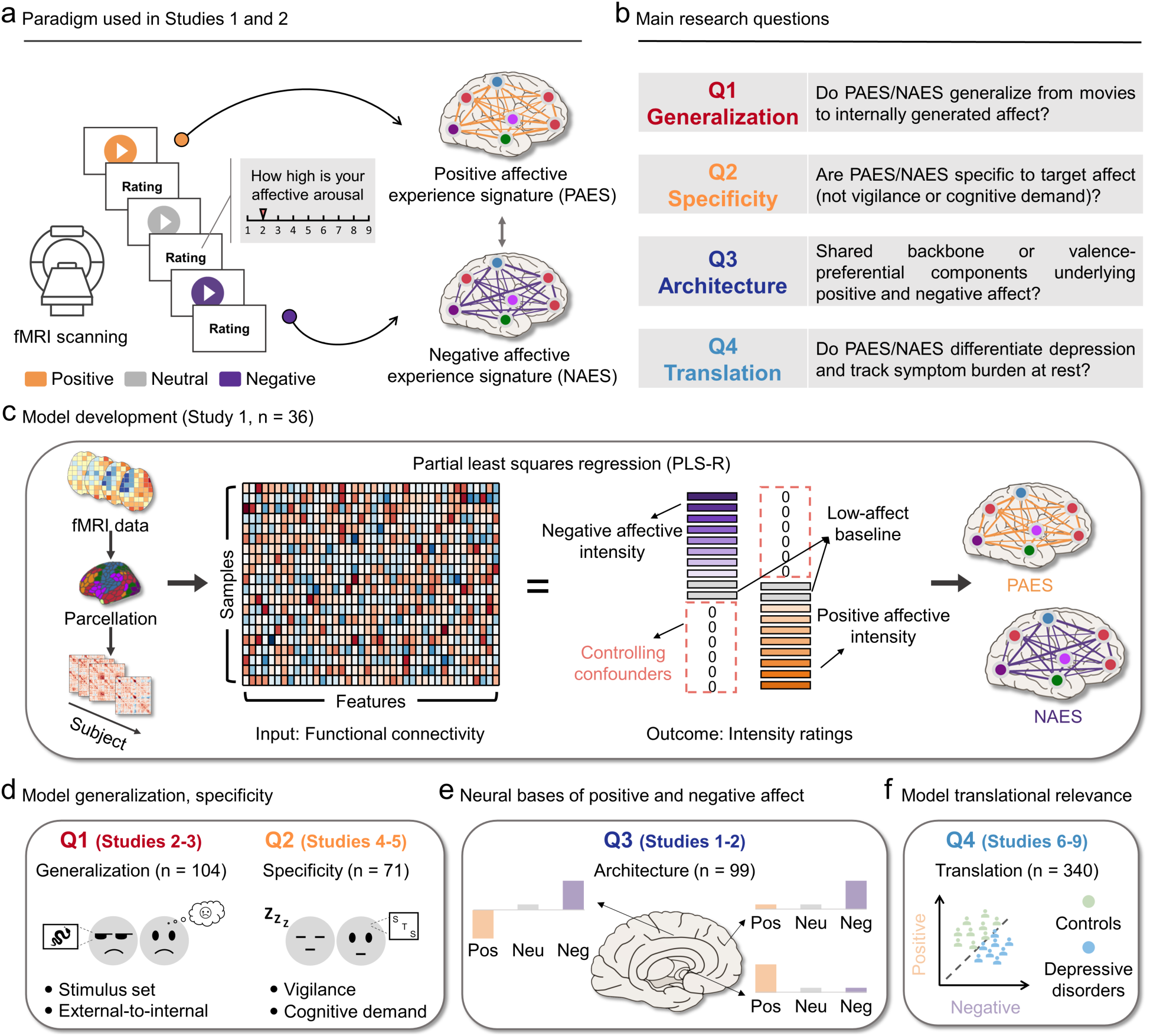
Overview of experimental paradigms, modeling strategy, and key research questions. **(a)**, Schematic of the experimental paradigm for Studies 1 and 2. Participants underwent fMRI scanning while viewing positive, neutral, and negative naturalistic movie clips, followed by self-reported ratings of affective intensity (i.e., affective arousal). These data were used to derive the Positive Affective Experience Signature (PAES) and Negative Affective Experience Signature (NAES). **(b)**, The four main research questions (Q1–Q4) addressed in this study, spanning model generalization, specificity, neural architecture, and clinical translation. **(c)**, Model development using partial least squares regression (PLS-R) in Study 1 (n = 36). Whole-brain functional connectivity (FC) features were used to predict positive and negative affective intensity. To enhance valence specificity, intensity ratings for the opposing valence condition were coded as zero (controlling confounders), yielding the PAES and NAES multivariate predictive patterns. **(d)**, Summary of the independent datasets used for model evaluation and application. Q1 was addressed by testing generalization to affect induced by a new stimulus set (Study 2, n = 63) and to internally generated affect (Study 3, n = 41). Q2 tested specificity against vigilance (sleep-wake state; Study 4, n = 28) and cognitive demand (working-memory load; Study 5, n = 43). **(e)**, Q3 aimed to reveal the shared and valence-preferential neural bases of positive and negative affective experience. **(f)**, Q4 evaluated clinical translation in resting-state cohorts of patients with depressive disorders versus healthy controls (Studies 6–9, n = 340).

## Results

### Development of whole-brain affective intensity signatures

To identify distributed neural representations of sustained positive and negative affective experience, we applied partial least squares regression (PLS-R) to whole-brain FC features from Study 1^7^ (n = 36) acquired during viewing of a curated set of 28 movie clips. Parcel-to-parcel FC, defined as Fisher z-transformed Pearson correlations between the time series of each pair of 468 parcels, was computed for each clip and participant (see Methods). These features were used to train models predicting valence-specific affective intensity (9-point Likert scale). Clips (∼70 s) were selected to elicit prolonged affective states and categorized as positive, neutral, or negative based on independent normative valence ratings and emotion-category assignments. An independent behavioral sample (n = 26) showed high agreement in valence (Intra-class Correlation Coefficient [ICC] = 0.99) and category labels (Fleiss’ κ = 0.83).

To model positive and negative affective intensity as partially independent, we used a two-channel outcome coding scheme (Fig. 1c). For PAES, intensity ratings for positive clips (range 2-9) were used as the target outcome and negative clips were coded as 0; for NAES, ratings for negative clips (range 2-9) were used and positive clips were coded as 0 (for a similar strategy, see ref. ^12^). Here, 0 serves as an off-target label for model estimation and is not a psychometric value on the 1-9 rating scale. To define a low-affect baseline, we retained (for each participant) only those normatively neutral clips that the participant rated at the minimum intensity (rating = 1). No positive or negative clips received a rating of 1. Opposite-valence trials were coded as 0 (rather than 1) to avoid conflating strongly affective off-target trials with the low-affect baseline, which could increase cross-valence leakage and reduce valence specificity. This scheme avoids imposing a single bipolar valence axis during training while preserving coarse valence structure by assigning intensity to the corresponding valence channel and treating the opposite valence as off-target, enabling empirical tests of sign-inverted versus valence-preferential organization. This procedure yielded two whole-brain functional connectivity signatures: the Positive Affective Experience Signature (PAES) and the Negative Affective Experience Signature (NAES) (Fig. 2a; see also Supplementary Figs. 1 and 2 for bootstrap-thresholded maps).

**Fig. 2.**
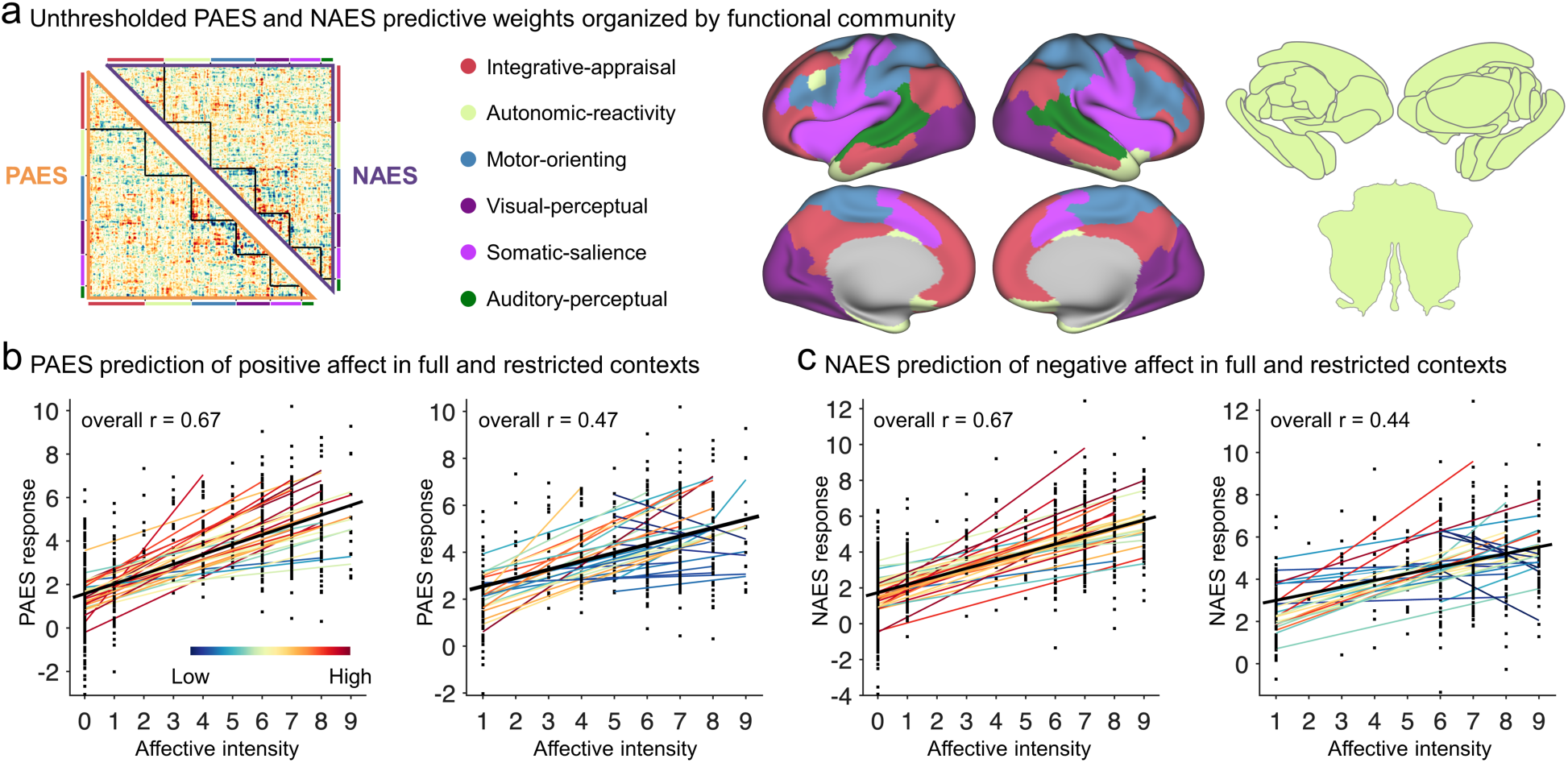
Whole-brain affective intensity signatures and predictive validity. **(a)**, Unthresholded Positive Affective Experience Signature (PAES) and Negative Affective Experience Signature (NAES), colored by community assignment to visualize the structured organization of the predictive patterns. A data-driven, bagging-enhanced Leiden community detection analysis identified six stable spatial communities, here termed the integrative-appraisal, autonomic-reactivity, motor-orienting, visual-perceptual, somatic-salience, and auditory-perceptual systems (see Results for details). **(b,c)**, Predictive validity of the signatures evaluated via 10-fold (participant-wise) cross-validation in the training cohort (Study 1, n = 36). Each colored line represents prediction within each individual participant. Black line indicates the overall (i.e., within and between participants) prediction. **(b)**, The PAES robustly predicted positive affective intensity (left panel: full context, i.e., evaluating across all clips with negative clips coded as 0; right panel: restricted context, i.e., excluding negative clips). **(c)**, The NAES robustly predicted negative affective intensity (left panel: full context, i.e., evaluating across all clips with positive clips coded as 0; right panel: restricted context, i.e., excluding positive clips). Both signatures maintained significant predictive performance even when opposite-valence trials were excluded, confirming they are not driven solely by a bipolar contrast.

### PAES and NAES robustly predict sustained positive and negative affect intensity

We next evaluated the predictive validity of the signatures. In 10-fold cross-validation within Study 1 (participant-wise folds), both signatures robustly tracked their target affective intensity (Figs. 2b, c). The PAES predicted positive experience intensity (with negative clips coded as 0; prediction–outcome correlation r = 0.67, 95% bootstrapped confidence interval [CI] = [0.63, 0.70], root mean square error [RMSE] = 2.36; both permutation P-values < 1 × 10^-4^). To ensure performance was not driven solely by contrasting opposing valences, we evaluated prediction after excluding negative clips. The PAES remained significantly predictive when restricted to positive and neutral trials (r = 0.47, 95% CI = [0.39, 0.54], RMSE = 2.60; both permutation P-values < 1 × 10^-4^). Similarly, the NAES predicted negative experience intensity (with positive clips coded as 0; r = 0.67, 95% CI = [0.63, 0.70], RMSE = 2.61; both permutation P-values< 1 × 10^-4^) and retained robust prediction when restricted to negative and neutral trials (r = 0.44, 95% CI = [0.36, 0.52], RMSE = 2.84; both permutation P-values < 1 × 10^-4^).

### Signatures generalize to an independent movie-viewing dataset

Having established the robust predictive validity of these signatures, we next examined their generalizability in an independent dataset with different participants and video stimuli (Study 2, n = 63)^7^, with no further model fitting. Participants in Study 2 viewed a different set of n = 40 naturalistic movie clips (lasting ∼25 s each) and provided post-clip affective intensity in the scanner (see Supplementary Materials). Given the relatively short duration of the clips, we concatenated within-condition time series across clips (positive, neutral, negative), yielding one FC map (and thus one signature expression) per condition and participant. Within-participant comparisons confirmed that all individuals reported higher affective intensity during positive and negative movie clips relative to neutral clips (Fig. 3a, left panel). As shown in the middle panel of Fig. 3a, using a two-alternative forced-choice (2AFC) classification framework (Methods), the PAES discriminated positive from negative conditions with 98.41 ± 1.58% standard error (SE) accuracy (two-tailed binomial test P < 1 × 10^-20^, area under the curve (AUC) = 0.999, Cohen’s d = 3.17) and positive from neutral with 98.41 ± 1.58% accuracy (P < 1 × 10^-20^, AUC = 0.998, d = 2.60). Conversely, the NAES discriminated negative from positive conditions with 98.41 ± 1.58% accuracy (P < 1 × 10^-20^, AUC = 0.997, d = 2.81) and negative from neutral with 96.83 ± 2.21% accuracy (P = 4.44 × 10^-16^, AUC = 0.991, d = 2.50; Fig. 3a, right panel). These large effects (d ≥ 2.50) indicate that both signatures generalize across independent participants and stimulus sets.

**Fig. 3.**
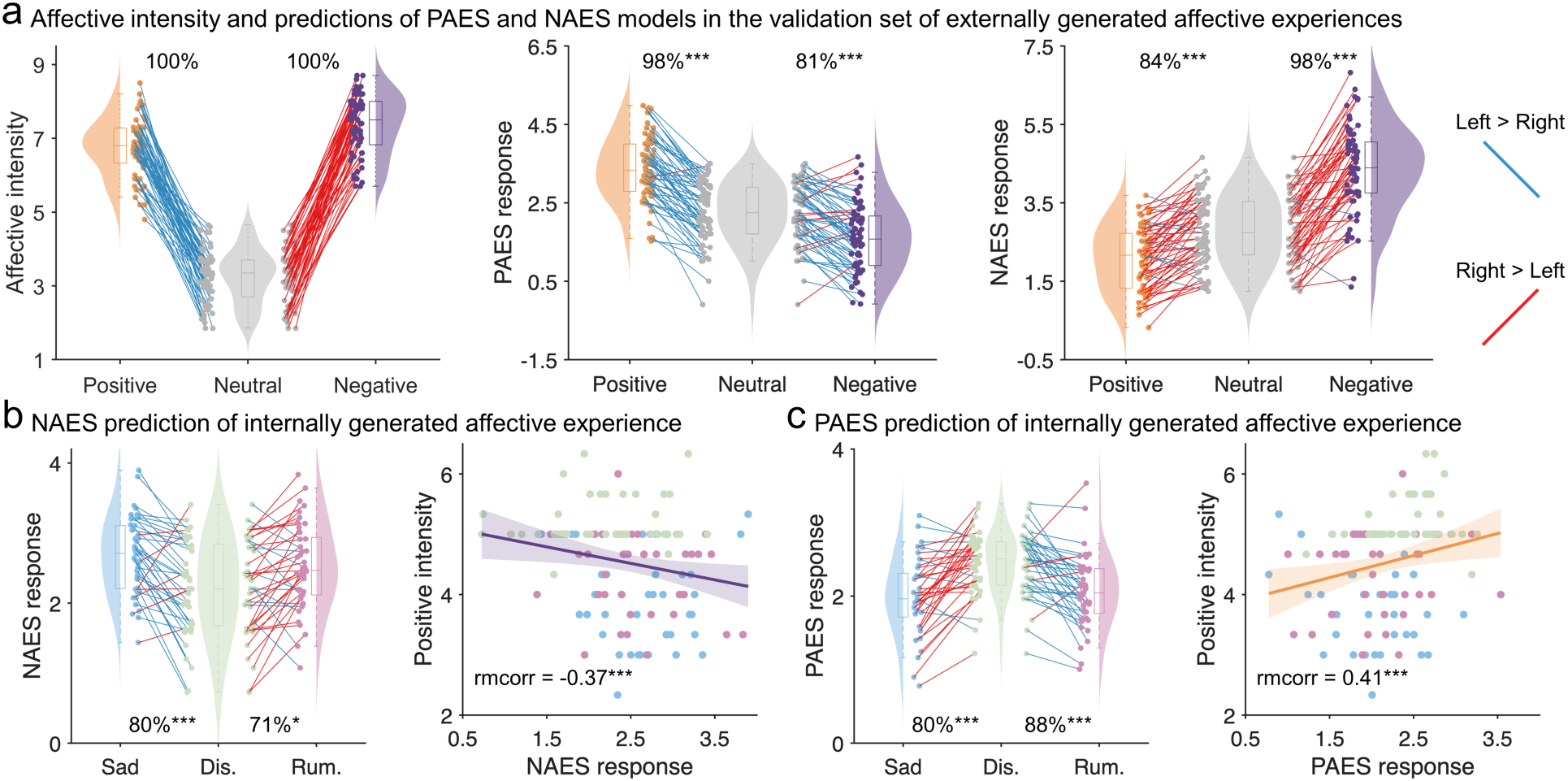
Generalization of PAES and NAES to independent external and internally generated affective states. **(a)**, Validation in an independent movie-viewing cohort (Study 2, n = 63). The left panel confirms that within-participant affective intensity was significantly higher for positive and negative clips relative to neutral clips. The middle and right panels show two-alternative forced-choice (2AFC) classification accuracies for PAES and NAES, respectively. Both signatures generalized to independent participants and stimuli, successfully discriminating target valences from neutral and opposite-valence conditions. **(b,c)**, Generalization to concept-driven, internally generated affect (Study 3, n = 41). Participants engaged in text-prompted sad memory recall, depressive rumination, and a neutral distraction control. **(b)**, NAES expression successfully distinguished internally generated negative states (sad memory and rumination) from the distraction control via 2AFC classification. Furthermore, NAES expression negatively covaried with subjective happiness ratings across conditions. **(c)**, PAES expression tracked the relative positivity of the distraction condition, successfully discriminating it from sad memory and rumination. PAES expression positively covaried with subjective happiness ratings across all internally generated states. For scatter plots in b and c, colored dots indicate different conditions (sad memory, distraction, and rumination), fitted lines depict the population-level estimate from a linear mixed-effects model, and shaded regions indicate the 95% confidence intervals. rmcorr, repeated measures correlation. * P < 0.05; ** P < 0.01; *** P < 0.001.

### Signatures accurately capture internally generated affect

We next tested whether signatures trained on externally elicited affect could generalize to internally generated states. To do so, we used an open dataset (Study 3, n = 41)^43^. Participants were presented with on-screen prompts designed to induce negative, self-referential processing, including sad memory recall (e.g., thinking about individual negative autobiographical events) and depressive rumination (e.g., analyzing their own personality to understand why they felt so depressed during the recently recalled events), alongside a neutral distraction control (imagining non-emotional scenarios).

Given that behavioral assessments in this open dataset were limited to a single “happiness” scale (1 = very unhappy to 9 = very happy), we used lower happiness scores as a proxy for negative affective intensity. Behaviorally, the distraction condition elicited higher happiness ratings than both memory recall and rumination^43^ (Supplementary Fig. 3), providing a behavioral benchmark for validation. Mirroring this subjective drop in positivity, NAES expression successfully distinguished the internally generated negative states from the distraction control (Fig. 3b, left panel). Specifically, using a 2AFC classification approach, NAES classified sad memory versus distraction with 80.49 ± 6.19% accuracy (P = 1.12 × 10^-4^, AUC = 0.88, d = 1.10) and rumination versus distraction with 70.73 ± 7.11% accuracy (P = 0.012, AUC = 0.79, d = 0.75). Beyond categorical discrimination, NAES expression covaried with subjective experience: repeated-measures correlation revealed a negative association with happiness ratings across all conditions (r_81_ = -0.37, P = 5.95 × 10^-4^, 95% CI = [-0.50, -0.23]; Fig. 3b, right panel).

Conversely, as expected based on the dimensional structure of affective experience, PAES expression tracked the relative positivity of the distraction condition (Fig. 3c, left panel), discriminating distraction from sad memory (80.49 ± 6.19% accuracy, P = 1.12 × 10^-4^, AUC = 0.90, d = 1.26) and from rumination (87.80 ± 5.11% accuracy, P = 7.84 × 10^-7^, AUC = 0.90, d = 1.11). PAES expression also covaried with subjective experience, showing a positive association with happiness ratings across conditions (r_81_ = 0.41, P = 1.07 × 10^-4^, 95% CI = [0.30, 0.51]; Fig. 3c, right panel). Together, these findings indicate that PAES and NAES capture core affect-related FC patterns that generalize from percept-driven (movie) to concept-driven (internally generated) experience.

### Signatures do not track vigilance or cognitive demand

As a further test of specificity, we evaluated whether PAES and NAES reflect valence-specific affective experience rather than broader valence-invariant state factors, focusing on two potential confounds: vigilance (sleep-wake state) and cognitive demand (working-memory load).

We first examined whether signature expression simply indexed sleep-wake state (i.e., fluctuations in vigilance/alertness) using an open dataset with concurrent fMRI and electroencephalogram (EEG) during wakeful rest and sleep (Study 4; n = 28)^44^. EEG-defined 30-s epochs were labeled as wakefulness or sleep. Participants contributed comparable numbers of wakefulness and sleep epochs (wakefulness: 56.36 ± 5.41; sleep: 40.21 ± 5.14; paired t-test t = -1.56, P = 0.131). Neither PAES nor NAES discriminated wakefulness from sleep using a 2AFC classification framework (PAES: 46 ± 9.43% accuracy, P = 0.85, AUC = 0.37, d = -0.38; NAES: 54 ± 9.43% accuracy, P = 0.85, AUC = 0.51, d = -0.04; Supplementary Fig. 4), indicating that signature expression is not driven by sleep-wake state.

We next tested whether PAES and NAES track cognitive demand using an open dataset with fMRI during spatial and verbal working memory tasks at high versus low load (Study 5; n = 44 participants were enrolled, with valid data for each task ranging from 35 to 43 participants)^45^. As expected, behavioral accuracy decreased as load increased (P-values < 5.99 × 10^-3^), confirming successful manipulation. However, neither PAES nor NAES significantly discriminated 2-back from 1-back conditions, nor either task condition from baseline, in a 2AFC framework (all P-values > 0.18 except the 2-back versus baseline contrast in spatial working memory task under the delayed large-reward condition (P = 0.09); see Supplementary Table 1). These findings indicate that PAES and NAES are not explained by simple working-memory load.

Together, these analyses provide little evidence that PAES or NAES are driven by vigilance or cognitive demand.

### PAES and NAES support a partially separable organization of sustained affect

Having found little evidence that PAES and NAES reflect broad valence-invariant state factors, we next asked whether sustained positive and negative affect are better captured as opposite poles of a single bipolar axis or as partially separable representations. To this end, we first examined cross-valence prediction in Study 1. PAES showed a weak negative association with negative affective intensity (excluding positive clips; r = -0.11, 95% CI = [-0.20, -0.01]; permutation P = 0.073), and NAES weakly negatively predicted positive affective intensity (excluding negative clips; r = -0.20, 95% CI = [-0.29, -0.11]; permutation P = 0.003). Cross-valence effects were substantially smaller than within-valence effects, arguing against a simple sign-inverted account. Moreover, the negative direction of cross-valence prediction is inconsistent with a valence-invariant intensity account, which would be expected to yield positive cross-valence associations. Together, these findings favor partially separable representations over a single bipolar axis expressed with opposite signs.

We next asked whether each signature showed the expected valence-specific ordering across conditions in Study 2: strong target-valence expression relative to neutral, accompanied by a weaker off-valence effect in the opposite direction. As shown in Fig. 3a, PAES expression was higher for neutral than negative condition, enabling discrimination of neutral versus negative conditions with 80.95 ± 4.95% accuracy (P = 7.46 × 10^-6^, AUC = 0.903, d = 1.32). Conversely, NAES expression was higher for neutral than positive condition, enabling discrimination of neutral versus positive conditions with 84.13 ± 4.60% accuracy (P = 3.38 × 10^-8^, AUC = 0.921, d = 1.41). This monotonic pattern is inconsistent with a purely valence-invariant intensity account and instead supports valence-specific organization of affective intensity representations. Consistent with the weak cross-valence predictions in Study 1, effect-size asymmetries in this independent dataset further argue against a simple sign-inverted account. Specifically, PAES showed approximately twofold larger separation for positive versus neutral than for neutral versus negative, and NAES showed approximately 1.8-fold larger separation for negative versus neutral than for neutral versus positive.

As a more explicit test of a bipolar valence axis account, we additionally trained a single support vector regression (SVR) model^5–7^ to predict signed valence ratings, constructed by recentering intensity ratings to 0-8 and multiplying negative-clip ratings by -1 (yielding -8 to +8, with 0 indicating neutral). Trained on Study 1 (cross-validated prediction-outcome r = 0.68; Supplementary Fig. 5a) and evaluated out-of-sample in Study 2 using the same within-subject 2AFC framework, this bipolar model reliably discriminated positive versus neutral and negative versus neutral conditions with similarly high accuracy (92.06 ± 3.41% [d = 2.09] and 88.89 ± 3.96% [d = 2.06], respectively; Supplementary Fig. 5b), consistent with a symmetric bipolar valence axis representation. However, its performance was lower than the corresponding target-valence versus neutral discrimination achieved by PAES (98%) and NAES (97%), indicating that separating positive and negative affective intensity into two valence-specific signatures provides additional sensitivity beyond a single bipolar axis model.

### Community-level characterization of PAES and NAES

To characterize the large-scale architecture of sustained affect, we assessed community structure in the combined PAES and NAES multivariate weight patterns using bagging-enhanced Leiden community detection^46^ (see Methods), and then asked how affective information was distributed within and between these communities. This analysis identified six stable communities (Bagged modularity Q = 0.57, mean ± SD Adjusted Rand Index [ARI] = 0.58 ± 0.07), corresponding to distinct functional systems supporting sustained affect. These comprised (1) an integrative-appraisal system anchoring the DMN (n = 74) and FPN (n = 35) networks, implicated in the construction of conscious affective experience^38^; (2) an autonomic-reactivity system comprising the vast majority of subcortical (STC; n = 36), cerebellar (CB; n = 32), and limbic (LN; n = 20) regions^47^; (3) a motor-orienting system integrating Somatomotor (SMN; n = 34) regions with the nearby dorsal attention network (DAN; e.g., frontal eye fields; n = 34), potentially supporting behavioral expressions such as approach and avoidance; (4) a visual-perceptual system dominated by the visual network (VN; n = 60); (5) a somatic-salience system (akin to the action-mode network) linking somatomotor (n = 36) regions with the ventral attention network (VAN; n = 26), implicating interoceptive processing and the homeostatic prioritization of internal states^48^; and (6) a small auditory-perceptual system located in the middle and superior temporal gyri (n = 12, 7, and 5 for DMN, SMN, and VAN, respectively). In Fig. 2a, regions are colored by these community assignments to visualize the structured, system-level organization of the predictive patterns.

To identify community-level connectivity patterns that were both independently predictive and important for intact model performance, we combined permutation-based sufficiency and importance analyses and retained only effects significant in both (FDR-corrected permutation P < 0.05; Fig. 4). This overlap revealed a distributed backbone shared by positive and negative affective experience. The integrative-appraisal system emerged as the principal common hub, showing shared predictive and performance-relevant coupling with the autonomic-reactivity and visual-perceptual systems, as well as within-system connectivity, consistent with a role in higher-order appraisal and affective integration. The somatic-salience system also emerged as a prominent connector at the between-system level, participating in shared interactions with the motor-orienting, visual-perceptual, and auditory-perceptual systems, consistent with a role in linking salience and interoceptive processing with sensory-affective information. In addition, connectivity within the motor-orienting and visual-perceptual systems, as well as between them, met both criteria for PAES and NAES, highlighting a prominent perceptual–orienting backbone. In the context of naturalistic movie viewing, this pattern is consistent with sustained affect being embedded in the transformation of external sensory input into orienting and action-related states, including approach- and avoidance-related tendencies. Together, these shared effects support a broadly distributed architecture that integrates appraisal, bodily-state, perceptual, and action-related processes, broadly consistent with appraisal-based, embodied, and constructionist views of emotion^49–51^.

**Fig. 4.**
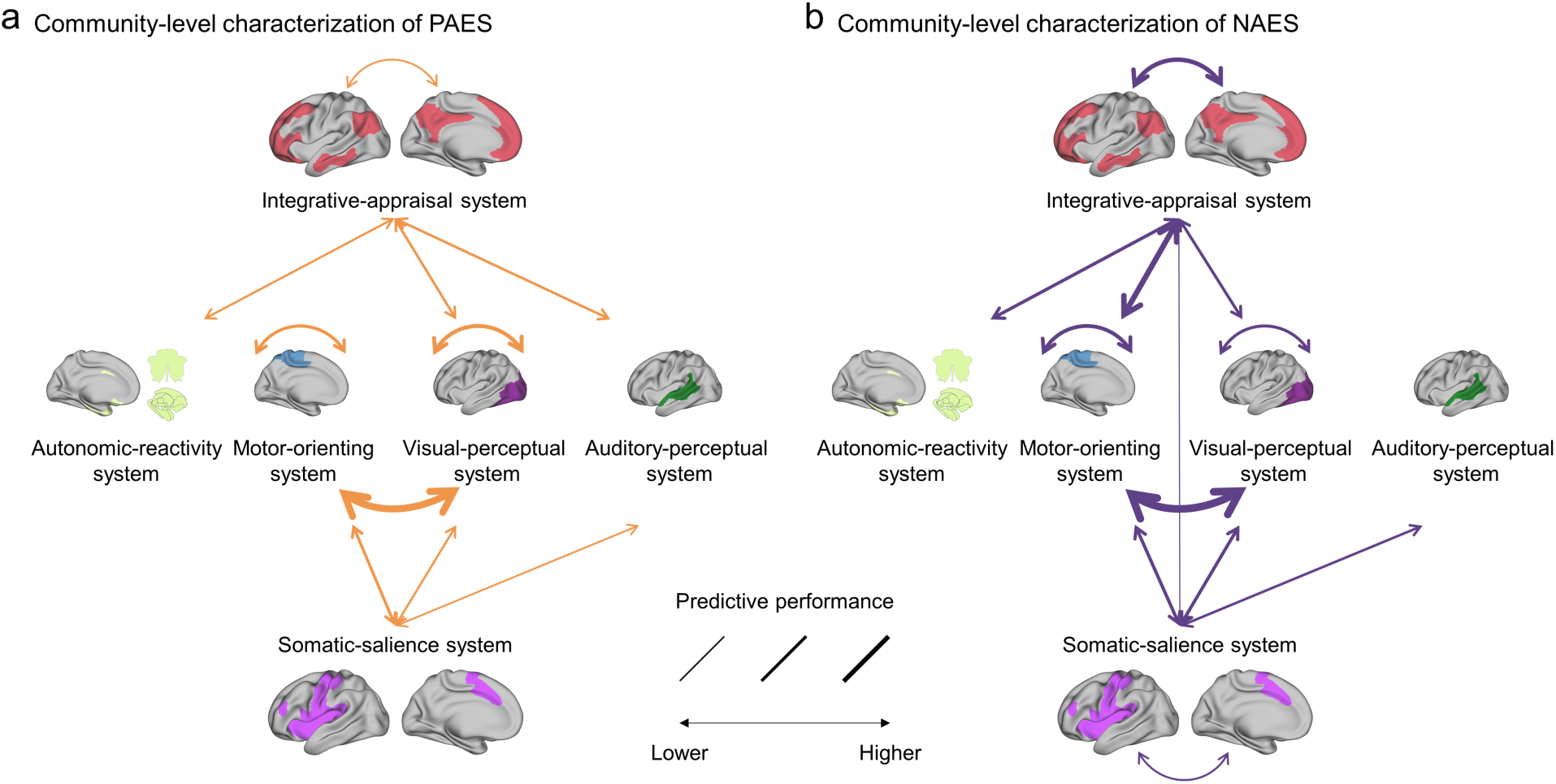
Sufficiency and importance of within- and between-community connectivity for PAES and NAES. **(a)**, Connectivity contributing to the Positive Affective Experience Signature (PAES). **(b)**, Connectivity contributing to the Negative Affective Experience Signature (NAES). Shown are within-community and between-community connectivity patterns that were identified as both sufficient and important for prediction. Sufficiency was assessed by evaluating the predictive performance of each pattern in isolation, whereas importance was assessed by permutation-based disruption of the corresponding connectivity pattern in the full model. Line thickness indicates predictive performance in the sufficiency analysis, with thicker lines denoting stronger predictive contribution.

Beyond this shared backbone, PAES additionally involved integrative-appraisal–auditory-perceptual connectivity, whereas NAES showed stronger involvement of the somatic-salience system, including within-system effects and additional coupling between the somatic-salience and integrative-appraisal systems. NAES also showed additional coupling between the integrative-appraisal and motor-orienting systems, a pattern that may reflect enhanced action-readiness or defensive orienting during negative affect. Together, these system-level findings point to a macroscale backbone shared by sustained positive and negative affective experience, alongside possible valence-preferential configurations.

### Region-level characterization of PAES and NAES

To characterize the PAES and NAES at a finer scale and identify more specific functional contributions, we conducted a parcel-wise local prediction analysis to estimate subjective affective experience. For each parcel, we tested whether its whole-brain connectivity profile alone could (i) predict affective intensity in Study 1 and (ii) discriminate affective conditions in Study 2.

This analysis identified multiple distributed regions whose connectivity profiles were individually predictive of sustained affective experience. As shown in Fig. 5a, for PAES, connectivity profiles in bilateral middle cingulate cortex (MCC), bilateral anterior insula (aIns), left anterior ventromedial prefrontal cortex, and left ventral striatum (ventromedial putamen) significantly predicted positive affective intensity in Study 1 (FDR-corrected permutation P < 0.05) and validated in Study 2 to discriminate positive from neutral and negative conditions (binomial test P < 0.05; higher expression for positive experience). For NAES, connectivity profiles in bilateral dorsomedial and dorsolateral prefrontal cortex (dmPFC, dlPFC), left MCC, left thalamus, bilateral caudal hippocampus, and anterior/posterior insula significantly predicted negative affective intensity in Study 1 (FDR-corrected permutation P < 0.05) and validated in Study 2 to discriminate negative from neutral and positive conditions (binomial test P < 0.05; higher expression for negative experience; Fig. 5b).

**Fig. 5.**
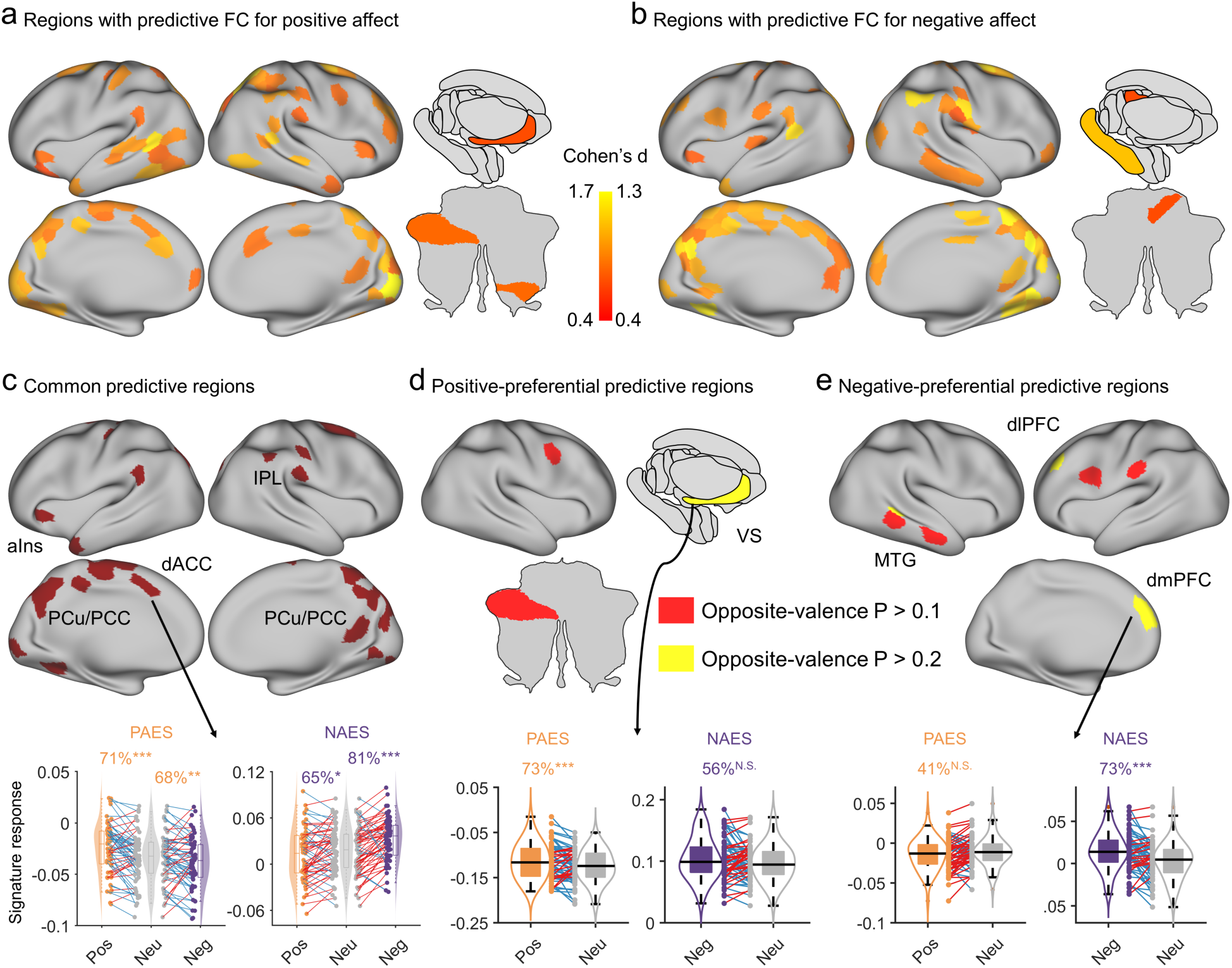
Parcel-wise local prediction analysis reveals shared substrates and valence-preferential double dissociations. **(a,b)**, Brain regions whose individual whole-brain connectivity profiles significantly predicted target affective intensity in Study 1 (FDR-corrected permutation P < 0.05) and were further validated by successfully discriminating corresponding affective conditions in Study 2 (binomial test P < 0.05). **(a)**, Regions predicting positive affective experience, including the bilateral middle cingulate cortex (MCC), bilateral anterior insula (aIns), left anterior ventromedial prefrontal cortex, and left ventromedial putamen. **(b)**, Regions predicting negative affective experience, including the bilateral dorsomedial and dorsolateral prefrontal cortex (dmPFC, dlPFC), left MCC, left thalamus, bilateral caudal hippocampus, and anterior/posterior insula. **(c)**, Shared neural substrates. Multiple regions, particularly within the somatic-salience system (e.g., MCC, aIns), integrative-appraisal system (e.g., precuneus [PCu], posterior cingulate cortex [PCC], inferior parietal lob [IPL]), and the visual-perceptual system, exhibited significant predictive profiles for both positive and negative affect. **(d,e)**, Double dissociation in valence-preferential sufficiency. **(d)**, Regions including the ventral striatum (VS) demonstrated selective sensitivity for positive affect, whereas their corresponding NAES profiles failed to predict negative outcomes in either Study 1 or 2. **(e)**, Conversely, regions including the dmPFC, dlPFC and middle temporal gyrus (MTG) demonstrated selective sensitivity for negative affect, whereas their corresponding PAES profiles failed to predict positive outcomes in either Study 1 or 2. To thoroughly illustrate this valence selectivity, thresholds for non-significant predictions in the opposite, non-target valence are explicitly denoted (displaying both P > 0.1 and P > 0.2). Box plots show parcel-based predictions for Study 2. Pos, positive affect; Neu, neutral affect; Neg, negative affect. *** P < 0.001; N.S., not significant.

Across signatures, several regions—particularly within the somatic-salience system (e.g., MCC and aIns), integrative-appraisal system (e.g., precuneus and posterior cingulate cortex [PCC]), and visual-perceptual system—showed significant predictions for both positive and negative affective intensity (Fig. 5c), consistent with partially shared substrates supporting sustained affective engagement (see also Supplementary Fig. 6 for a signed overlap map of bootstrap-stable PAES and NAES features). Importantly, within this shared distributed backbone, we observed a striking double dissociation in valence-preferential sufficiency (Figs. 5d, e). For example, the ventral striatum showed selective sensitivity to positive affect: its connectivity profile predicted positive experience in Study 1 and discriminated positive from neutral in Study 2, whereas the corresponding NAES profile did not predict negative outcomes (P > 0.2). Conversely, left dmPFC and dlPFC showed selective sensitivity to negative affect: their connectivity profiles predicted negative experience in Study 1 and discriminated negative states in Study 2, whereas the corresponding PAES profiles did not predict positive outcomes (P > 0.2). This pattern is consistent with preferential recruitment of cortico-striatal valuation-related circuitry for positive affective intensity and greater engagement of dorsomedial/lateral prefrontal systems involved in appraisal and emotion regulation-related processing during negative affective states.

Finally, a region-based virtual lesion analysis indicated that removing the connectivity of any single parcel produced only minimal performance degradation (e.g., prediction performance change < 0.75% for PAES and < 0.53% for NAES). This indicates that both signatures reflect distributed, redundant representations rather than relying on a small set of localized hubs.

### Signatures capture affective dysregulation in resting-state depression cohorts

A critical next question was whether PAES and NAES extend from externally evoked and internally generated affective states observed under laboratory conditions to spontaneous affective dysregulation in clinical populations. Because depressive disorders are characterized by protracted negative affect and anhedonia, even in the absence of explicit cognitive tasks, we hypothesized that signature expression during stimulus-free resting-state fMRI (rs-fMRI) would index baseline affective burden and differentiate patients from healthy controls in the predicted opposing directions: specifically, depressive disorder samples would show reduced PAES expression and increased NAES expression relative to controls. To test this hypothesis, we applied the signatures to four independent resting-state datasets comprising first-episode, unmedicated adults and adolescents with depressive disorders, alongside healthy controls.

Across three independent cohorts of first-episode, unmedicated adults with major depressive disorders (MDD; Studies 6–8; total N = 214), both signatures successfully classified patients versus controls (see Fig. 6a–d, left panels, for Studies 6 and 7). Specifically, PAES classification accuracies ranged from 65.35% to 69.57%, while NAES accuracies ranged from 68.32% to 71.74% (Table 1). Furthermore, in cohorts with available clinical measures (Studies 6 and 7), signature expression dimensionally tracked symptom burden (Fig. 6a–d, right panels). In Study 6^52^, PAES was negatively associated with Beck Depression Inventory (BDI) scores across all participants (r_64_ = -0.29, P = 0.019, 95% CI = [-0.50, -0.04]) and NAES was positively associated (r_64_ = 0.37, P = 0.002, 95% CI = [0.12, 0.56]). Within the MDD group alone, NAES remained significantly associated with BDI (r_31_ = 0.36, P = 0.042, 95% CI = [0.06, 0.65]), whereas PAES showed a trend-level negative association (r_31_ = -0.34, P = 0.052, 95% CI = [-0.65, 0.002]). These symptom associations showed consistent directional trends in Study 7 (measured via Hamilton Depression Rating Scale (HAMD); PAES: r_20_ = -0.38, P = 0.080, 95% CI = [-0.66, 0.11]; NAES: r_20_ = 0.40, P = 0.064, 95% CI = [-0.07, 0.74]), though limited statistical power in this smaller sample precluded strict two-tailed significance.

**Fig. 6.**
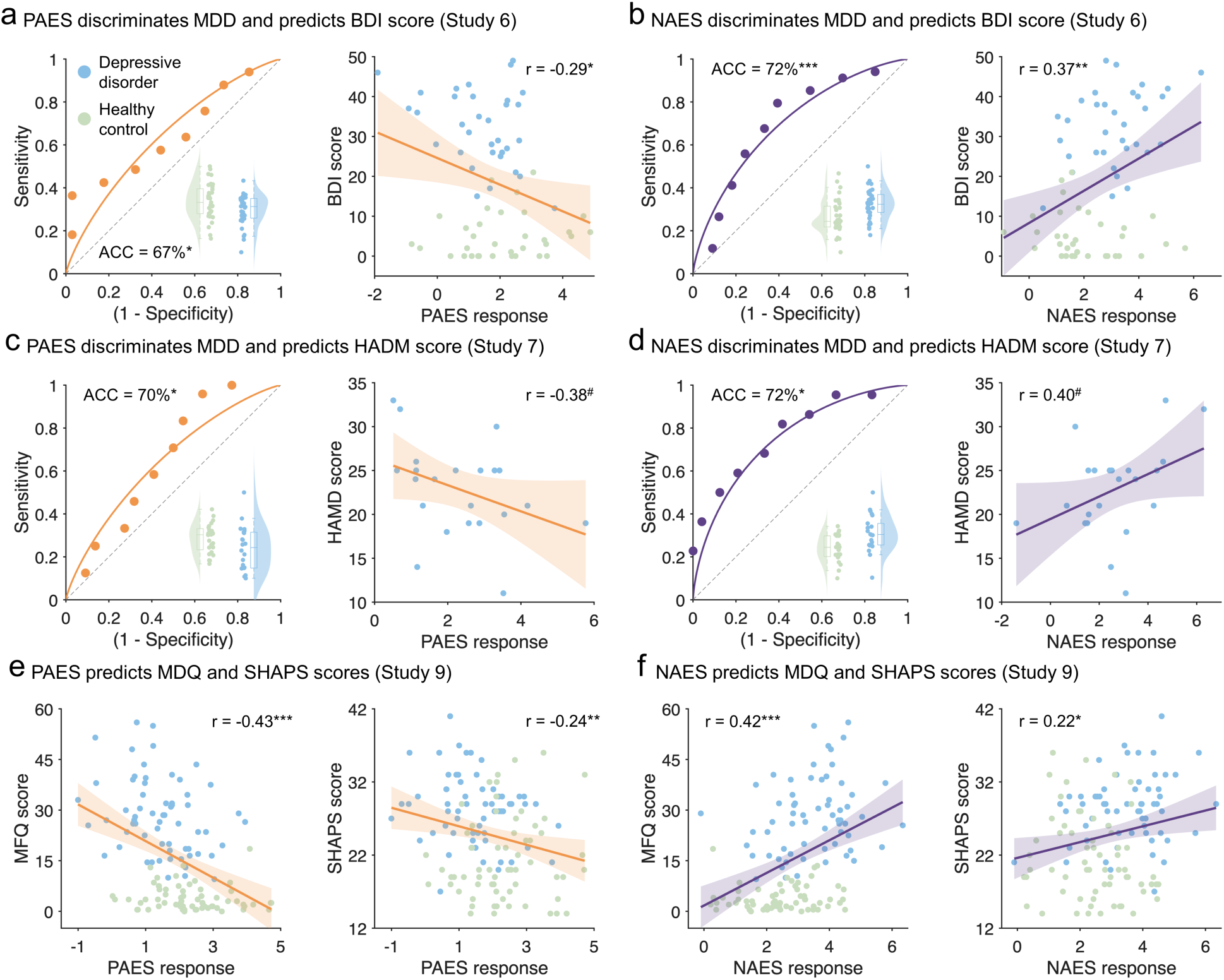
Clinical generalization and symptom tracking of PAES and NAES in independent resting-state depression cohorts. **(a–d)**, Case–control discrimination and symptom tracking across two independent resting-state fMRI datasets (Study 6, n = 67; Study 7, n = 46). Left panels: violin plots display PAES and NAES expression in healthy controls (HC) and patients with major depressive disorders (MDD). Patients exhibit significantly reduced PAES expression and elevated NAES expression. Receiver operating characteristic (ROC) curves show classification performance between MDD and HC. Right panels: signature expression tracks general depression severity. PAES negatively correlates, and NAES positively correlates, with Beck Depression Inventory (BDI) scores in Study 6 and Hamilton Depression Rating Scale (HAMD) scores in Study 7. **(e,f)**, Signature expression maps onto specific symptom dimensions in the adolescent cohort (Study 9, n = 126). PAES **(e)** negatively correlates, while NAES **(f)** positively correlates, with overall mood symptoms (Mood and Feelings Questionnaire [MFQ]) and anhedonia (Snaith-Hamilton Pleasure Scale [SHAPS]). For all scatter plots, solid lines depict the linear fit and shaded regions indicate the 95% confidence intervals. ^#^ P < 0.1; * P < 0.05; ** P < 0.01; *** P < 0.001.

**Table 1.** PAES and NAES classify depressive disorders versus controls.

|  | PAES | NAES |
| --- | --- | --- |
| Study 6<br>MDD: n = 34;<br>HC: n = 33 | ACC = $67.16 \pm 5.74\%$<br>$P = 9.66 \times 10^{-3}$<br>AUC = 0.64<br>d = 0.52 | ACC = $71.64 \pm 5.51\%$<br>$P = 8.11 \times 10^{-4}$<br>AUC = 0.73<br>d = 0.76 |
| Study 7<br>MDD: n = 22;<br>HC: n = 24 | ACC = $69.57 \pm 6.78\%$<br>$P = 0.025$<br>AUC = 0.67<br>d = 0.54 | ACC = $71.74 \pm 6.64\%$<br>$P = 0.011$<br>AUC = 0.79<br>d = 0.97 |
| Study 8<br>MDD: n = 52<br>HC: n = 49 | ACC = $65.35 \pm 4.74\%$<br>$P = 6.76 \times 10^{-3}$<br>AUC = 0.66<br>d = 0.49 | ACC = $68.32 \pm 4.63\%$<br>$P = 8.91 \times 10^{-4}$<br>AUC = 0.69<br>d = 0.66 |
| Study 9<br>Depression: n = 63<br>HC: n = 63) | ACC = $69.05 \pm 4.12\%$<br>$P = 2.28 \times 10^{-5}$<br>AUC = 0.72<br>d = 0.82 | ACC = $69.05 \pm 4.12\%$<br>$P = 2.28 \times 10^{-5}$<br>AUC = 0.74<br>d = 0.89 |

PAES, positive affective experience signature; NAES, negative affective experience signature; ACC, accuracy (mean ± standard error [SE]); AUC, area under the receiver operating characteristic curve; d, Cohen's d; MDD, major depressive disorders; HC, healthy controls.

Highlighting the developmental generalization of these markers, both signatures also significantly distinguished adolescents with depressive disorders from healthy controls (Study 9^53^; N = 126; Table 1). Moreover, signature expression mapped onto specific symptom dimensions: PAES expression was negatively correlated with overall mood symptoms (Mood and Feelings Questionnaire [MFQ]; r_124_ = -0.43, P = 5.90 × 10^-7^, 95% CI = [-0.55, -0.28]) and anhedonia (Snaith-Hamilton Pleasure Scale [SHAPS]; r_123_ = -0.24, P = 0.008, 95% CI = [-0.39, -0.06]; Fig. 6e), whereas NAES expression was positively correlated with these measures (MFQ: r_124_ = 0.42, P = 9.70 × 10^-7^, 95% CI = [0.28, 0.53]; SHAPS: r_123_ = 0.22, P = 0.013, 95% CI = [0.05, 0.38]; Fig. 6f). These dimensional relationships were largely maintained within the adolescent depressive disorder group alone, further linking PAES to positive affect deficits and NAES to elevated negative affect. Specifically, PAES expression was negatively correlated with MFQ (r_61_ = -0.29, P = 0.019, 95% CI = [-0.48, -0.09]) and SHAPS (PAES r_60_ = -0.22, P = 0.081, 95% CI = [-0.42, -0.001]), while NAES expression showed positive associations with MFQ (r_61_ = 0.22, P = 0.082, 95% CI = [0.02, 0.41]) and SHAPS (r_60_ = 0.22, P = 0.080, 95% CI = [-0.02, 0.45]).

Together, PAES and NAES consistently differentiated depressive disorder groups from healthy controls across four independent resting-state datasets and showed convergent associations with symptom burden and anhedonia across distinct clinical instruments (BDI, HAMD, MFQ, SHAPS). Notably, depressive phenotypes and symptom scales are heterogeneous, and these effects emerged in task-free resting-state fMRI in which participants’ ongoing experience is unconstrained; these factors may reduce observed brain–symptom associations. Overall, the cross-cohort replication and dimensional symptom tracking provide evidence that PAES and NAES capture clinically relevant variation in task-free resting-state connectivity.

### Resting-state functional connectivity-based case-control decoders show modest performance and limited cross-cohort generalization

To contextualize the clinical generalization of PAES and NAES, we trained support vector machine (SVM)-based rs-fMRI case-control classifiers separately in each depression cohort (Studies 6–9) and evaluated their performance both within the training cohort using leave-one-out cross-validation and across the other cohorts.

As shown in Table 2, these cohort-specific decoders yielded only modest performance within their respective cohorts and generalized inconsistently across datasets. This pattern may reflect heterogeneity in resting-state functional connectivity and depressive phenotypes, and is consistent with prior evidence that larger samples are needed to derive stable dysfunction-related markers. By contrast, the observation that PAES and NAES generalized across multiple independent resting-state depression cohorts supports the possibility that experimentally derived affective signatures may provide a more portable index of clinically relevant affective dysfunction.

**Table 2.** Within- and cross-cohort performance of cohort-specific rs-fMRI depression decoders. Each decoder was evaluated within its training dataset using leave-one-out cross-validation and then tested on the remaining independent cohorts without further refitting. Shown are classification accuracy (ACC), two-sided binomial P value, area under the receiver operating characteristic curve (AUC), and Cohen’s d.

| Development | Test |  |  |  |
| --- | --- | --- | --- | --- |
|  | Study 6 | Study 7 | Study 8 | Study 9 |
| Study 6 (n = 67) | ACC = 62.69% | ACC = 52.17% | ACC = 53.47% | ACC = 52.38% |
|  | P = 0.066 | P = 1.000 | P = 0.766 | P = 0.656 |
|  | AUC = 0.64 | AUC = 0.48 | AUC = 0.46 | AUC = 0.46 |
|  | d = 0.31 | d = -0.19 | d = -0.16 | d = -0.13 |
| Study 7 (n = 46) | ACC = 53.73% | ACC = 73.91% | ACC = 64.36% | ACC = 59.52% |
|  | P = 0.715 | P = 0.004 | P = 0.012 | P = 0.040 |
|  | AUC = 0.46 | AUC = 0.85 | AUC = 0.66 | AUC = 0.60 |
|  | d = -0.17 | d = 1.3 | d = 0.47 | d = 0.34 |
| Study 8 (n = 101) | ACC = 56.72% | ACC = 69.57% | ACC = 61.39% | ACC = 58.73% |
|  | P = 0.393 | P = 0.025 | P = 0.058 | P = 0.061 |
|  | AUC = 0.54 | AUC = 0.70 | AUC = 0.69 | AUC = 0.57 |
|  | d = 0.04 | d = 0.74 | d = 0.67 | d = 0.26 |
| Study 9 (n = 126) | ACC = 58.21% | ACC = 65.22% | ACC = 68.32% | ACC = 59.52% |
| | P = 0.271 | P = 0.103 | P = $8.90 \times 10^{-4}$ | P = 0.040 |
|  | AUC = 0.51 | AUC = 0.66 | AUC = 0.69 | AUC = 0.64 |
|  | d = -0.12 | d = 0.59 | d = 0.58 | d = 0.38 |

## DISCUSSION

Our findings suggest that sustained affect is supported by a distributed neural architecture that bridges externally elicited and internally generated experience. Using whole-brain functional connectivity, we identified positive and negative affective experience signatures (PAES and NAES) whose expression tracked valence-specific affective intensity, generalized across independent externally evoked and internally generated affective states, and showed little evidence of being explained by vigilance (sleep-wake state) or cognitive demand (working-memory load) under the conditions tested. Together with the asymmetry in predictive performance, systems- and region-level characterization supported a shared distributed backbone and valence-preferential components, consistent with a partially separable organization of sustained positive and negative affect. Extending beyond experimentally evoked states, the signatures also differentiated depression cohorts from healthy controls in opposing directions and tracked symptom burden and anhedonia, supporting their relevance to task-free affective dysregulation.

By modeling within-valence intensity and treating opposite-valence trials as off-target outcomes, we minimized reliance on a simple positive–negative contrast or a single bipolar valence axis. Both PAES and NAES retained robust performance when evaluation was restricted to within-valence trials, whereas cross-valence associations were weak and negative across datasets, arguing that the signatures capture valence-specific sustained affective intensity rather than merely reflecting a unitary affective-intensity axis. Because whole-brain FC can be sensitive to broad state factors^54^ we tested signature specificity in independent datasets indexing vigilance and cognitive engagement; signature expression showed little evidence of tracking vigilance or cognitive demand under the conditions tested. Collectively, these results indicate that PAES and NAES capture valence-specific sustained affective intensity rather than valence-invariant intensity or non-affective state confounds, and support a partially separable rather than strictly bipolar account of sustained affective experience.

Given that the signatures are defined on whole-brain FC aggregated over extended epochs, they are well positioned to index sustained, state-like coordination among appraisal, memory, and interoceptive systems rather than fast, stimulus-locked sensory transients. This interpretation is consistent with the prominent contribution of a DMN-centered integrative-appraisal system^55^, which supports internally oriented cognition and the construction of affective meaning^56,57^. Because these systems are engaged during both externally elicited narratives^58^ and internally generated affect^59^, we reasoned that signatures derived from percept-driven affect might generalize to concept-driven affective states. Consistent with this view, signature expression differentiated internally generated negative states from neutral distraction and aligned with subjective ratings, supporting a shared connectivity substrate for sustained affect across external and internal modes of experience. At the same time, this generalization does not imply that externally elicited and internally generated affect are neurally identical; rather, it suggests partially shared large-scale representations that bridge these modes of experience. This generalization raises the possibility that the same signatures capture stimulus-free affective burden during resting-state fMRI.

Across four independent resting-state datasets, PAES and NAES distinguished depressive disorder groups from healthy controls in opposing directions (reduced PAES and elevated NAES) and tracked symptom burden and anhedonia. These effects in task-free resting-state fMRI align with depression as a disorder marked by persistent, stimulus-independent affect^20,21^. Although unconstrained resting-state fMRI cannot identify the content of ongoing thought, it offers high ecological validity for indexing stimulus-free, spontaneous affective processes that are known to be persistently altered in depressive disorders^60,61^. While case–control discrimination was moderate, this is expected given clinical heterogeneity and cross-site fMRI variability. Moreover, direct case–control decoders trained on resting-state functional connectivity within individual depression cohorts showed only modest within-cohort performance and limited cross-cohort generalization, consistent with recent work reporting similarly modest performance at comparable sample sizes^23^. Against this background, the ability of PAES and NAES to generalize across multiple independent resting-state depression cohorts suggests that experimentally anchored, theory-guided signatures may capture clinically relevant affective variation more portably than de novo diagnosis-trained resting-state decoders. Importantly, the convergence across cohorts and symptom dimensions supports the clinical relevance of signature expression. Beyond case–control discrimination, PAES and NAES provide quantitative indices conceptually aligned with the Research Domain Criteria (RDoC) Positive and Negative Valence Systems^62^ and may prove useful as dimensional markers of symptom burden and anhedonia, as well as candidate measures for future longitudinal and interventional studies. More broadly, these results link experimentally elicited sustained affect to clinically relevant resting-state variation.

Our findings also support a distributed functional architecture for sustained positive and negative affect, comprising both shared and valence-preferential components. At the community level, combining sufficiency and importance analyses identified a macroscale backbone shared by positive and negative affective experience. Within this backbone, the integrative-appraisal system emerged as the principal common hub, showing shared within-system effects and shared coupling with autonomic-reactivity and visual-perceptual systems, consistent with a role in higher-order appraisal and integration of affective experience^38,63,64^ and broadly compatible with appraisal theories of emotion^50^. The somatic-salience system additionally emerged as a prominent connector, linking motor-orienting, visual-perceptual, and auditory-perceptual systems, suggesting that sustained affect engages salience- and interoception-related processes alongside sensory-affective processing^65^, in a manner consistent with constructionist views emphasizing bodily-state and interoceptive contributions to emotional experience^51^. Shared effects within and between the motor-orienting and visual-perceptual systems further suggest that sustained affect in naturalistic settings is embedded in ongoing perceptual processing and motor-orienting dynamics^37,66^, aligning with embodied views that situate emotion within perception-action couplings rather than purely abstract representations^49^. This distributed organization aligns with prior multivariate decoding work across affective domains^5,7,10–14,67^ and argues against a localized “affect center” ^68,69^. At the regional level, multiple parcels carried informative connectivity profiles for both positive and negative affective intensity, including salience regions (e.g., MCC and aIns), integrative-appraisal regions (e.g., PCu/PCC/IPL), and visual cortex, suggesting shared substrates for sustained affective engagement and meaning construction. Within this distributed backbone, we also observed valence-preferential profiles: ventromedial putamen connectivity was preferentially informative for positive affect, whereas dmPFC/dlPFC connectivity was preferentially informative for negative affect, consistent with differential contributions of valuation-related versus appraisal/control-related circuitry^56,70,71^. In movie-viewing datasets where outcomes were explicitly valence-specific, weaker cross-valence than within-valence prediction, together with this regional double dissociation, supports a partially separable organization of sustained affect rather than a strictly bipolar implementation. Although these valence-preferential interpretations are supported by validation in an independent dataset, they should be interpreted with the caveat that non-target tests were not significant and do not by themselves establish equivalence. Applications to internally generated affect and resting-state clinical cohorts address generalization and clinical relevance, but do not provide a symmetric test of common-pattern versus partially separable organization.

Several limitations motivate future work. First, to minimize in-scanner fatigue, we modeled within-valence sustained intensity by combining post-clip affective intensity with normatively validated valence labels. Although independent behavioral validation showed high agreement, future work incorporating continuous, within-individual valence ratings could more directly dissociate valence and intensity dynamics. Second, the internally generated affect dataset provided an important test of cross-mode generalization, but its single happiness rating did not permit a fully symmetric assessment of positive and negative affective intensity. Third, although PAES and NAES generalized across multiple clinical cohorts, case–control discrimination for depressive disorders was moderate, potentially reflecting clinical heterogeneity and the pooling of heterogeneous affective categories within each valence during model development. Signatures optimized for more specific affective states may improve sensitivity to particular clinical phenotypes or symptom dimensions (e.g., sadness- or joy-focused signatures relevant to depression). Finally, our depression cohorts primarily comprised first-episode, unmedicated adults and adolescents, reducing treatment-related confounds but leaving open questions about generalizability to chronic or treatment-resistant populations and about how signature expression evolves with pharmacological or neuromodulatory interventions.

In sum, our findings reveal a distributed neural architecture of sustained affect that generalizes across externally elicited and internally generated experience and extends to task-free clinical contexts. By delineating shared and valence-preferential components of sustained affect, PAES and NAES provide a framework for quantifying affective experience and its dysregulation in psychiatric disorders.

## Methods

Data from nine independent fMRI studies were used to develop, validate, and test out-of-sample FC-based models of within-valence affect intensity. Study 1 served as the training dataset, Studies 2–3 tested generalization to independent movie stimuli and internally generated affect, Studies 4–5 assessed specificity against sleep-wake state and cognitive load, and Studies 6–9 evaluated clinical generalization in depression cohorts. Key MRI data acquisition parameters are provided in Supplementary Table 2. MRI data were preprocessed using fMRIPrep 22.0.2^72^ to correct for head motion and spatially normalize images to a common neuroanatomical space (MNI152NLin6Asym), followed by smoothing (6 mm full-width at half maximum), nuisance regression, and bandpass filtering (see Supplementary Materials for details).

The research protocols were approved by the Southwest University ethics committee (No. H24037) and adhered to the latest Declaration of Helsinki. Written informed consent was obtained from all participants before participation. For publicly available datasets, data collection was approved by the respective local institutional review boards of the original studies, and all participants (or their legal guardians) provided written informed consent prior to data collection. For all other independent datasets, detailed ethics approvals are reported alongside the specific cohort descriptions.

### Participants in Study 1

Thirty-eight healthy, right-handed students from the University of Electronic Science and Technology of China (UESTC) were recruited to undergo fMRI while performing a movie-based subjective affect rating task. Following our previous study^7^, two participants with abnormally low affective intensity to emotional videos during scanning (all ratings < 3) were excluded before any analyses, resulting in a final sample of 36 participants (17 females; mean ± SD age = 21.03 ± 2.22 years). All participants reported no current or past physical, neurological, or psychiatric disorders and no MRI contraindications. Written informed consent was obtained before participation. The study protocol was approved by the UESTC ethics committee (No. 06142292724710) and adhered to the latest Declaration of Helsinki.

### Stimulus selection and normative validation in Study 1

Candidate movie stimuli were selected from a large pool of clips sourced from publicly available online media and screened by two of the authors (R.Z. and F.Z.). A total of 28 movie clips were selected for the fMRI study based on behavioral ratings obtained from an independent sample (n = 26). Participants rated each clip on 9-point Likert scales for affective intensity (i.e., affective arousal; 1 = very low, 9 = very high), valence (1 = very negative, 5 = neutral, 9 = very positive), and emotional consistency (1 = very low, 9 = very high; consistency of emotional experience across the clip duration), and indicated the dominant emotional experience evoked by each clip (fearful, disgusting, sad, ambiguous negative, happy, exciting, romantic, ambiguous positive, or neutral). When multiple positive (or negative) emotions were endorsed for a given clip, the label ambiguous positive (or ambiguous negative) was assigned. Collapsing categorical labels into positive, negative, and neutral valence categories, mean valence ratings were 7.81, 2.34, and 5.00, respectively. For inclusion, positive clips were required to have mean normative valence > 7, negative clips mean normative valence < 3, and neutral clips mean normative valence within a narrow midpoint range (4.5-5.5). Inter-rater reliability for valence ratings was excellent (ICC = 0.99). We quantified inter-rater agreement in categorical assignment using Fleiss’ κ. When clips were grouped into positive (happy, exciting, romantic, ambiguous positive), negative (fearful, disgusting, sad, ambiguous negative), and neutral categories, agreement across the 28 clips was very high by conventional benchmarks (Fleiss’ κ = 0.83). All clips showed high within-clip emotional consistency (mean consistency > 7.2).

### Affective intensity rating paradigm in Study 1

Participants viewed 28 videos across three runs: 9 positive, 9 negative, and 10 neutral clips. Each run included three or four clips from each category. The sequence of runs and trials was fixed across participants, with the condition order pseudorandomized within this fixed sequence. Each video lasted ∼70 s, followed by a fixation cross (1-1.2 s). Participants then reported their affective intensity (i.e., affective arousal) during the preceding clip using a 9-point Likert scale (1 = extremely low, 9 = extremely high) during a 5 s response window, followed by a 10-12 s washout period. Stimulus presentation and behavioral data collection were controlled using MATLAB 2014a (Mathworks, Natick, MA) and Psychtoolbox (http://psychtoolbox.org/).

### Brain parcellations

The parcellation used to develop connectivity-based affective experience signatures included 400 cortical regions from the Schaefer Atlas^73^, 36 subcortical regions from the Brainnetome Atlas^74^ and 32 cerebellar regions from the SUIT Atlas^75^ (total = 468 parcels).

### Participant-specific trial inclusion for model development

We applied participant-specific filtering to reduce contamination of the low-affect baseline by idiosyncratic affective responses. Specifically, for each participant we retained only neutral clips (as defined by normative valence and category labels) that the participant rated with the minimum affective intensity (rating = 1), thereby restricting the low-affect baseline to minimal-intensity trials at the individual level. Conversely, we confirmed that no positive or negative clips had rating 1. These criteria were intended to improve affective category specificity in the training data.

### Brain model development and evaluation

We used PLS-R to develop valence-specific models that predict affective intensity from brain connectivity during positive and negative affective experiences. PLS-R estimates two sets of latent variables: brain components (FC maps) and affective intensity factors, optimized to maximize their intercorrelation (i.e., maximizing the explanation of variance in ratings with brain patterns). Unlike standard single-outcome approaches, PLS-R jointly estimates two outcomes, yielding separate brain patterns for positive- and negative-affective outcomes and enabling simultaneous prediction of correlated affective states^12^.

We first derived affect-related FC features for each participant. To account for hemodynamic lag, we shifted the fMRI time series by approximately 4 s relative to the video onsets. Next, we computed parcel-to-parcel FC (Fisher z-transformed Pearson correlation coefficients) during each video clip viewing, for each participant separately. These video-level FC features were then submitted to a PLS-R framework implemented via MATLAB’s ‘plsregress’ function. Predictors (X) comprised whole-brain FC maps stacked across participants. The outcome (Y) matrix contained affective intensity for each valence condition in separate columns (Y1 for positive, Y2 for negative). Ratings for the non-target valence were set to 0 in the corresponding column, constraining each solution to be valence-specific. The resulting brain patterns predictive of Y1 and Y2 constitute models optimized for positive and negative affective intensity, respectively. Each model was projected as a single predictive spatial map.

To evaluate how well each PLS-R brain pattern (i.e., PAES and NAES) predicted targeted affective experiences (and to probe specificity), we used 10-fold cross-validation in Study 1. Folds were defined at the participant level to prevent leakage across clips from the same individual. To quantify accuracy and model fit, we report both prediction-outcome Pearson correlation and RMSE.

### Evaluating the generalizability of PAES and NAES

We applied the signatures to an independent validation cohort (Study 2, n = 63; age = 19.87 ± 2.01, 25 females)^7^ to obtain signature responses for each map, defined as the dot product between the pattern weight map and the test FC map. Study 2 used a similar fMRI paradigm but entirely different video stimuli in an independent sample (see Supplementary Materials for details), providing a stringent external validation and an out-of-sample assessment of sensitivity and specificity. Experimental procedures of Study 2 were approved by the local ethics committee at UESTC (No.06142292724710), in accordance with the latest version of the Declaration of Helsinki. Classification performance was assessed with 2AFC tests within individuals by comparing signature responses for two conditions and selecting the condition with higher PAES (or NAES) expression as the target-valence condition. P values were calculated using a two-sided binomial test. Notably, we did not exclude any trials in this dataset. We conducted three comparisons: positive versus neutral, negative versus neutral, and positive versus negative.

To additionally evaluate the predictive capacity of PAES and NAES in the absence of external stimuli, we tested model responses in a publicly available dataset (Study 3, n = 41; age = 22.7± 4.1; 22 females)^43^. Participants completed a modified rumination-state task during fMRI, engaging in three prompt-guided mental states: sad memory (e.g., thinking about individual negative autobiographical events), rumination (e.g., analyzing their own personality to understand why they felt so depressed in the events they just remembered), and distraction (e.g., thinking about the layout of a typical classroom). Before rumination and distraction, participants generated keywords to cue the recall of negative life events, inducing a sad mood and facilitating entry into rumination (see Supplementary Materials for details). Behavioral results indicated that distraction involved greater imagery, was more positive, and increased happiness relative to rumination and sad-memory states (see Chen et al.^43^). Accordingly, we assessed the classification accuracy of the two models for internally generated affect using within-subject forced-choice tests, including sad memory versus distraction and rumination versus distraction as well as associations between signature responses and happiness ratings after each mental state. Participants completed the same paradigm three times using different MRI scanners. We averaged functional connectivity and subjective ratings for each condition across these three sessions.

### Evaluating PAES and NAES specificity

We assessed model specificity using two independent open datasets: a simultaneous fMRI-EEG sleep-wake dataset (Study 4)^44^ and a cognitive load (n-back) fMRI dataset (Study 5)^45^.

For Study 4, we analyzed 28 participants (from an initial sample of 33; age = 22.1 ± 3.2 years; 16 females) who completed two resting-state and two sleep sessions and exhibited both sleep-wake state (EEG-labeled in the original dataset). We calculated the mean PAES and NAES responses for wake and sleep conditions for each subject and performed 2AFC classification tests to determine if the signatures could discriminate between states.

Study 5 included typically developing children (n = 43; age = 10.4 ± 0.96 years; 14 females) selected from a larger cohort that also included children with Attention-Deficit/Hyperactivity Disorder (ADHD). Participants performed eight n-back working memory tasks in a block design, varying by domain (verbal and spatial), load (baseline, 1-back and 2-back), reward (small and large) and feedback time (immediate and delayed); for full details of the task design and behavioral procedures, see ref. ^45^. In the verbal task, participants indicated whether the current letter matched the letter presented n steps earlier. In the spatial task, they indicated whether the current letter’s position matched the position presented n steps earlier. The tasks included three block types: 1-back (48 trials), 2-back (48 trials), and baseline (12 trials). Each task run consisted of one 1-back block, one 2-back block, and two baseline blocks. Participants were told that they would receive either small ($0.02) or large ($0.25) rewards for correct responses. To minimize potential trial-by-trial affective responses to feedback, we restricted our analysis to tasks with delayed feedback (where participants viewed a fixation cross between trials and received performance summaries only at the end of blocks), excluding those with immediate feedback. We derived condition-specific FC for each participant; to account for hemodynamic lag, condition onsets were shifted by approximately 4 s. We performed a total of 12 2AFC comparisons to test whether the signatures tracked cognitive load. These comparisons contrasted load conditions (2-back versus 1-back, 2-back versus baseline, and 1-back versus baseline) separately for each domain (verbal, spatial) and reward level (small, large).

### Community detection and network architecture

To characterize the stable functional architecture of the predictive signatures, we performed a bagging-enhanced community detection analysis designed to ensure robustness to sampling variability. Instead of clustering a single fixed weight map, we generated 10,000 bootstrap realizations of the predictive model by resampling clip-level FC observations (participant × clip) with replacement and re-training the PLS regression for each iteration. For each bootstrap iteration, we combined the positive (PAES) and negative (NAES) weight matrices to define a functional connectivity profile for each of the 468 brain regions.

We then constructed a weighted graph for each iteration using a k-nearest neighbor (k-NN) approach^76^. We calculated the Euclidean distance between regional predictive profiles and identified the k nearest neighbors for each region. To determine the optimal neighborhood size (k), we employed a two-stage stability optimization procedure. First, we performed a coarse sweep of candidate values (k in [5, 10, 15, 20, 25, 30, 40]), which identified k=10 as the preliminary optimum based on a combined score of modularity (Q) and solution stability (ARI). The modularity score (Q) quantifies the density of connections within communities relative to a random null model, serving as a measure of network definition, while the ARI measures the consistency of cluster assignments across bootstrap iterations (0 = random, 1 = perfect consistency). Second, we performed a fine-grained search around this value (k in [6, 7, …, 14]), identifying k = 9 as the global optimum that maximized the stability-modularity trade-off.

For each bootstrap iteration, edges in the graph were weighted by their Hadamard overlap—a measure of the proportion of shared neighbors—to prioritize topological structure over simple magnitude. We then applied the Leiden community detection algorithm to each of the 10,000 graphs to partition the regions into communities. To derive a final consensus partition, we computed a stability matrix encoding the probability that any pair of regions was assigned to the same community across these 10,000 bootstrap partitions. Finally, we applied the Leiden algorithm to this stability matrix, yielding a robust consensus partition of the brain into six stable functional systems.

### Community-level importance analysis

To quantify the contribution of community-level connectivity patterns to model performance, we conducted a permutation-based blockwise disruption analysis. Connectivity blocks were defined as edges within a community or between a pair of communities. For each block, the corresponding features were permuted in the held-out test data within each cross-validation fold, while the trained model was kept fixed. The same permutation was applied to all features in a block, disrupting the association between that block-specific connectivity pattern and the outcome while preserving the covariance structure within the block. Predictions were aggregated across folds, and performance was quantified separately for PAES and NAES as the pooled Pearson correlation between predicted and observed values across all held-out observations. This procedure was repeated 10,000 times per block. Block importance was indexed as the decrease in prediction performance relative to the unpermuted baseline. Statistical significance was determined using a one-sided permutation test with FDR correction across blocks.

### Clinical datasets and classification analyses

Studies 6–9 were resting-state fMRI datasets comprising both patients with depressive disorders and healthy controls, acquired across different sites using diverse scanners. Detailed demographic information and key MRI acquisition parameters are provided in the Supplementary Materials. For each resting-state fMRI run, pattern expression values were calculated and submitted to a single-interval classification analysis to discriminate patients from controls. Classification thresholds were selected to maximize overall accuracy. Statistical significance of classification accuracy was assessed using two-sided binomial tests. Area under the curve (AUC) is reported as a threshold-independent measure of classification performance.

### Support vector regression and support vector machine models

In line with our previous studies^5,7,67,77^, we used a linear SVR model with cost parameter C = 1 and epsilon = 0.1 for the bipolar valence analysis, and a linear SVM with cost parameter C = 1 for the rs-fMRI depression classifiers. These models were implemented using the Spider toolbox (http://people.kyb.tuebingen.mpg.de/spider).

## Acknowledgments

We thank all researchers who collected and/or shared the datasets included in this work. This work was supported by the National Natural Science Foundation of China (grant nos. 32571268 and 32300862) and the Chongqing Social Science Planning Project (grant no. 2024YC035).

## Author contributions

Conceptualization, B.B. and F.Z.; Investigation, R.Z., D.D., X.G., T.X., and B.C.; Formal analysis, R.Z., D.D., and F.Z.; Writing – original draft, R.Z., D.D., T.F., B.B., and F.Z.; Writing – review & editing, all authors; Funding acquisition, F.Z.; Supervision, T.F., B.B., and F.Z.

